# Coalescent models for populations with time-varying population sizes and arbitrary offspring distributions

**DOI:** 10.1101/131730

**Authors:** Christophe Fraser, Lucy M Li

## Abstract

The coalescent has been used to infer from gene genealogies the population dynamics of biological systems, such as the prevalence of an infectious disease. The offspring distribution affects the relationship between population dynamics and the genealogy, and for infectious diseases, the offspring distribution is often highly overdispersed. Here, we provide a general formula for the coalescent rate for populations with time-varying sizes and any offspring distribution. The formula is valid in the same large population limit as Kingman’s original derivation. By relating our derivation to existing formulations of the coalescent, we show that differences in the coalescent rate derived for many population models may be explained by differences in the offspring distribution. The coalescent derivations presented here could be used to quantify the overdispersion in the offspring distribution of infectious diseases, which is useful for accurate modelling disease outbreaks.

## 1 Introduction

Gene genealogies provide much useful information about the populations from which the genes are sampled, in particular how the population size changed over time in the past. This has proven especially useful in cases where other indicators of population size are limited, such as the study of extinct species over geological time, or the study of infectious disease outbreaks, where surveillance may be poor, or biased. The basic insight underlying this work is provided by the coalescent, which describes the statistical relationship between population size and the genealogy [1]. This statistical model can be fit to genealogical data to infer past population sizes. For a most simple example, the probability that two randomly chosen individuals are siblings is the inverse of the population size at the time when their parents were breeding. With time going backwards from the present, a coalescent model describes the changing probability over time that lineages from a sample coalesce, which occurs when two lineages share a common parent and correspond to branching events in the genealogy.

Kingman’s original coalescent model was formulated for a population reproducing in discrete generations with a constant population size, and for which only a small proportion of the population is sampled [1]. Since then, variants of the coalescent have been developed. These include models with overlapping generations evolving in continuous time [2], models with deterministically changing population size [3], fluctuating population sizes [4], heterogeneities in the offspring distribution [5], large sample fraction [6], population structure [7], stochastic population dynamics, or even multiple genes evolving through recombination and mutation [8]. There are many different methods for parameterising the inferred population sizes, which are parameters of the coalescent models. The approach used in [9] was to treat the population size as constant between each inter-coalescent time interval, resulting in a skyline plot. Many approaches have been used to smooth these plots [10]. An alternative approach is to use a parametric function, such an exponential function, as in [3], other smooth or piecewise curves, or where appropriate, epidemic models, which can be deterministic [11], or stochastic [12].

Despite the simple assumptions of Kingman’s coalescent, it is robust to changes in many of its assumptions. For example, it still holds true in the presence of fluctuations in population dynamics and fast migration rates between subpopulations [4]. However, the relationship between coalescent rate and the population size no longer hold when assumptions of the offspring distribution are violated. When the variance in the offspring distribution is greater than that of a Poisson distribution, the effective population size inferred using the coalescent is smaller than the actual population size. Kingman [1] showed that with a constant sized population, the ratio between the actual and effective population sizes is equal to the variance in the offspring distribution. This relationship might not hold in biological systems where the population size varies over time, such as during infectious disease epidemics. Because gene genealogies of pathogens are increasingly being used to infer properties of the epidemic, accurate formulation of the coalescent for time-varying populations with heterogeneous offspring distributions would make inference results from gene genealogies more accurate [13]. Previous derivations of the coalescent with time-varying population size and variable offspring distribution have shown that the above result is valid to describe the mean coalescent rate if the population size is replaced by its harmonic mean [14], however a derivation which allows time-varying coalescent rates is needed to infer time-varying population sizes use the coalescent.

Here, we present a variant on Kingman’s coalescent for the case of a population that is changing systematically over time and where individuals are heterogeneous in terms of the number of offspring they generate. We start with a very simple derivation for the case of discrete generations, analogous to Kingman’s derivation, and present a partial generalisation to continuous time. We show that in special cases, the simple formula is equal to other formulas derived under different assumptions, suggesting that our formula has generality beyond its restrictive assumptions, and that variance in the offspring distribution may be a key driver of differences between published models. We then derive the formula again for a related model in continuous time. The validity of the derivations have been demonstrated in a recently published work with simulated disease outbreaks [15].

An alternative framework for relating population dynamics to gene genealogies is the birth-death model (cite Stadler), which in our parlance corresponds to a geometric offspring distribution. Generalising birth-death models to other offspring distributions, or relating the two conceptually separate frameworks, is beyond the scope of this paper.

## 2 Theory and Results

### 2.1 Definition of the effective population size

We define the effective population size *N*_*e*_(*t*) by first defining the coalescent rate *p*_2_(*t*). *p*_2_(*t*)*dt* is the probability that any two selected lineages coalesce in the time interval [*t, t + dt*], where, as usual in coalescent models, time *t* goes backwards from the present. The coalescent rate is given by,

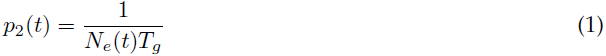

where *T*_*g*_ is the generation time. The overall coalescent rate is given by 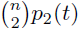, where 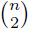 is the number of different pairs that could coalesce amongst a studied sample of size *n*. In the classical Fisher-Wright model, the population size is constant, denoted *N*, reproduction occurs in discrete generations of duration T_*g*_. Furthermore, all individuals have equal probability of producing offspring. More specifically, the number of offspring of a set of individuals in one generation is given by the multinomial with *N* probabilities = 1*/N*. When *N* is large, this is well approximated by a Poisson distribution with mean 1. Going back one generation, the probability that an individual shares a parent with any other individual is 1*/N*, and thus the coalescent probability is 1*/N* per generation, or approximately 1*/*(*NT*_*g*_) per unit of time. As expected, *N* = *N*_*e*_(*t*) for this classical Fisher-Wright coalescent.

### 2.2 Discrete generation model with arbitrary offspring distribution and changing population size

Instead of a Poisson distribution with mean 1, we consider the case of an arbitrary offspring distribution (Figure 1). To calculate the coalescent rate, we only need to specify the mean, commonly known as the reproduction number *R*(*t*), and the variance, which we denote *σ*^2^. The classical Fisher-Wright coalescent is recovered when *R*(*t*) = σ^2^ = 1. We denote the probability mass function of the offspring distribution *Φ*, such that Φ(ν) is the probability of an individual having *ν* offspring. It is normalised such that 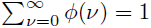 The mean of *Φ* is 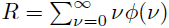 and its variance is 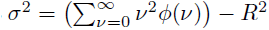.

**Figure 1.**
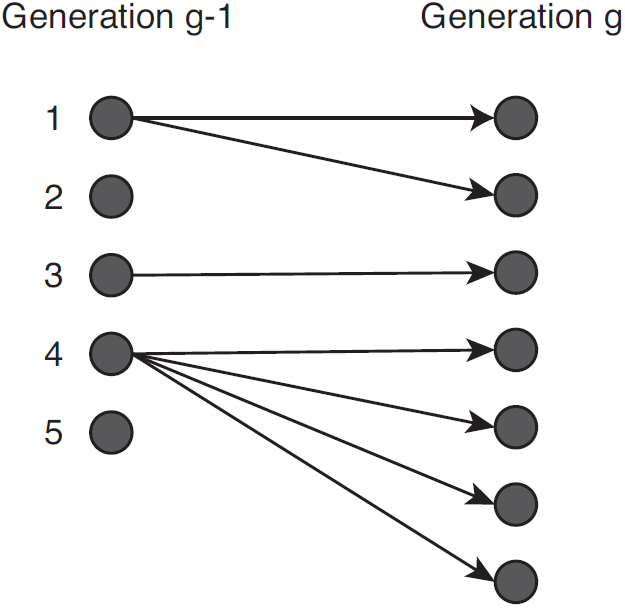
An illustrative example of a population undergoing one generation of reproduction, from *g* – 1 to *g*. *N*_*g*-1_ = 5 individuals in generation *g* – 1 are labelled 1,…, 5, and they have *v*_1_ = 2; *v*_2_ = 0; *v*_3_=1, *v*_4_ = 4 and *v*_5_ = 0 offspring, respectively. At generation *g*, the population size is *N*_*g*_ = *v*_1_ +…+ *v* _5_ = 7. The reproduction number *R*_*g*–1_ is the mean of *v*_i_, i.e. 7/5, which is also the relative change in population size *N*_*g*_/*N*_*g*–1_ The variance of the offspring distribution is V_g–1_ = ((v_1_ – *R*_*g*–1_)^2^ +…(v_5_– *R*_*g*-1_)^2^)/(5 –1) = 14/5. This is much greater than the value 7/5 expected under a homogeneous offspring model, indicating that the offspring distribution is skew. Two randomly selected members of generation g share the same parent if they are one of seven possible pairs, the offspring of 1 or any two of the four offspring of individual 4. There are 21 possible pairs, and so the coalescent probability is 7/21 = 1/3. The formula (V_g–1_/R_g–1_ +R_g–1_ –1)/Ng gives 24/70 = 0.34 which is very close to the exact result; the approximation rapidly converges to the exact result for large population sizes. This formula compares with the conventional approximation, 1/*H*(*N*) = 0.17, where *H* is the harmonic mean. The reason for the improvement is better representation of the heterogeneity in number of offspring and of the effects of changing population size.

Now consider a population evolving one generation from its ancestor at time *t - T*_*g*_ to time *t*. Label *i* = 1 *…N* (*t - T*_*g*_) all the individuals in the parent generation. Let *ν*_*i*_ be the number of each of their offspring, distributed according to *Φ*, i.e. *ν*_*i*_ *∼* Φ; then the number of offspring is 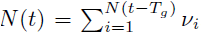. The total number of pairs in the population at time *t* is *N* (*t*)(*N* (*t*) - 1)*/*2, and the number of pairs that are siblings (i.e. share a parent) is 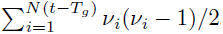. The probability of an arbitrarily chosen pair of isolates chosen at time *t* coalescing in the previous generation is the ratio of these quantities

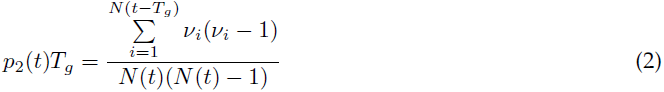

For a large population, the mean of the number of offspring is the mean of the offspring distribution, i.e.

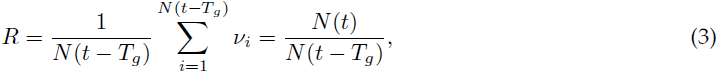

and the variance is the variance of the offspring distribution, i.e.

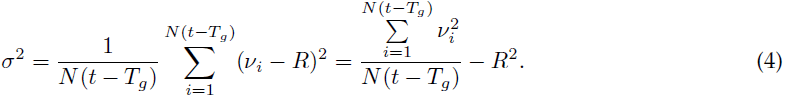

Inserting these equations back into Equation 2,

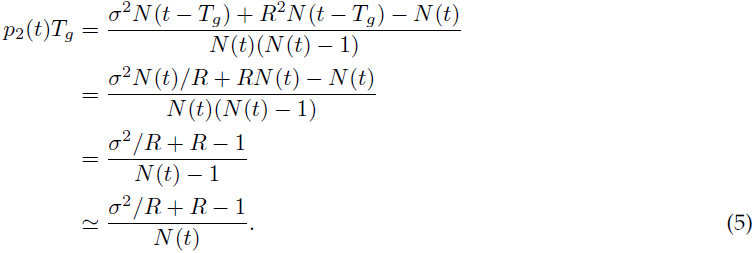

Substituting Equation 5 into Equation 1, we obtain this equation for the effective population size

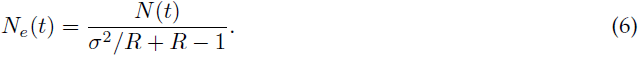

In Equations 5 and 6, the reproduction number and variance in offspring distribution stay constant over time. However, these can be replaced by time-varying values *R*(*t*) and *σ*(*t*) to allow temporal changes in the offspring distribution.

In the following sections (2.3-2.7), we show that Equation 6 reduces to some commonly used formulations of the coalescent.

### 2.3 Constant population size

In the case of constant population size (in which case *R* = 1), this reduces to the well-known equation originally proposed by Kingman [1]

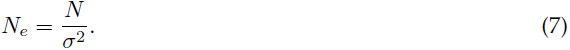

### 2.4 Poisson offspring distribution

In the case of Poisson offspring distribution (in which case *σ*^2^ = *R*), this reduces to

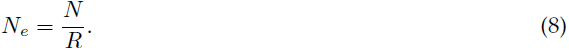

When the population size is constant as well (*R* = 1), then *N*_*e*_ = *N*, which corresponds to Kingman’s coalescent.

### 2.5 Geometric offspring distribution

In the case of a geometric offspring distribution *σ*^2^ = *R* + *R*^2^, so Equation 6 reduces to

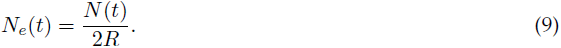

Using conventional parameterization of the geometric distribution, the mean *R* = (1 *- p*)*/p* refers to the expected number of success (offspring) given a probability of failure (not producing any offspring) of *p*. For the special case of constant sized population size with geometric offspring distribution, so that mean *R* = 1, Equation 9 reduces to the equation for the Moran model [1],

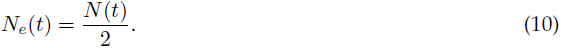

### 2.6 Negative binomial offspring distribution

A more flexible distribution is the negative binomial, which has an additional dispersion parameter *k* that controls the shape of the distribution. In the case of negative binomial distribution,

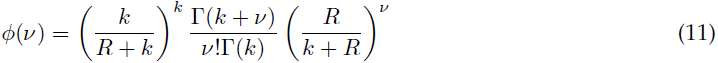

which has mean *R* and variance *σ*^2^ = *R* + *R*^2^/k, then

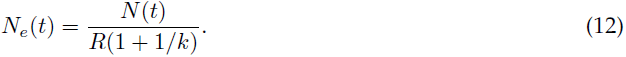

### 2.7 Exponential growth models

The exponential growth model is often used to capture the early stages of infectious disease spread in the population, during which the population size can change rapidly within a short period of time. Fitting the exponential model to pathogen genealogies can help to estimate the rate of epidemic growth [16].

Consider the special case when *N* (*t*_0_) = 1 at some initial time, then with discrete generation growth, *N* (*t*) = exp(*r*(*t - t*_0_)) = *R*^(*t-t*^0)*/T*_*g*_ with *r* = log(*R*)*/T*_*g*_, the exponential rate of growth. The effective population size is

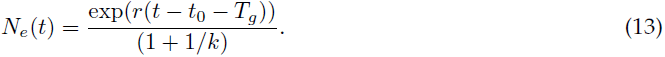

### 2.8 Continuous time models

Although the derivations in Section 2.2 were based on discrete generations, they have good agreement with results for continuous-time models, such as those derived by Volz et al. [11] for epidemic models. For example, the SIR model describes the changes over time of susceptible, infectious, and recovered individuals in a population:

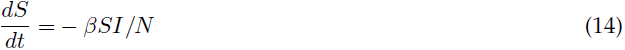

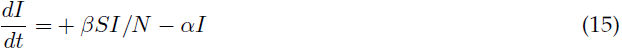

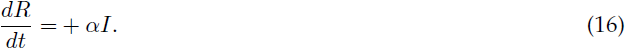

In the SIR model, the population size is *I*, the effective reproduction number is *βS/α*, and the generation time is *T*_*g*_ = 1*/α*. The offspring distribution is geometric because the number of transmissions per time unit is Poisson distributed and the duration of infectiousness is exponentially distributed. The effective population size is

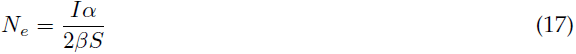

and the coalescent rate is

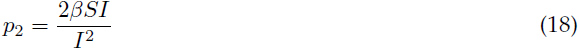

which is the formula derived by Volz et al. [11].

## 3 Discussion

Here we presented a flexible framework to model the coalescent process with any arbitrary offspring distribution. Kingman [1] showed that in a constant size population, the effective and actual population sizes are related via the variance in the offspring distribution. The formulation of the coalescent here demonstrates that the relationship depends both on the variance and the mean of the offspring distribution.

We also showed that many existing formulations of the coalescent were special cases of the general expression for coalescent rate presented here. This indicates that differences in the coalescent rate between these models are due to differences in the offspring distribution.

Our derivations for the negative binomial agree with those derived by Koelle and Rasmussen [17] in a constant sized population. However, our framework is more general in that it can be used to infer time-varying population sizes and any arbitrary offspring distribution.

Explicitly defining the coalescent rate in terms of both the population size and the offspring distribution is especially important for studying infectious disease spread, because overdispersion in the offspring distribution is a common phenomenon observed across many diseases [18]. Superspreading events are more likely when the offspring distribution is overdispersed, and this increases the uncertainty in predicting future epidemic trajectory and impacts the effectiveness of control strategies. Li et al. [15] used the derivations presented here and demonstrated that the variance of the offspring distribution could be estimated by fitting epidemic models to pathogen genealogy. In particular, Li et al. [15] demonstrate how to incorporate the derivations here into a practical skyline-type likelihood for parameter inference, both conventionally backwards in time, and also forwards in time as needed to fit dynamic epidemic models.

Although our derivations were based on population models with discrete generations, the agreement with the overlapping-generations Moran model and the SIR model [11] suggest that our derivations could approximate some continuous time models. However, the approximations would likely not work for infectious diseases with longer generation times, during which the coalescent rate could significantly change.

In particular, we did not attempt derivations for more complex epidemiological models such as the SEIR model, which present the additional challenge that the number infected and infectious individuals are no longer equal [17] and the ratio between the two may change over time during an epidemic. We show in separate work that the correspondence becomes more complicated and less accurate than that explored here [15].

Just as in Kingman’s coalescent, we assumed that no more than one coalescent event occurs at the same time. However, when the variance in the offspring distribution is high, multiple mergers could occur in the gene genealogy of the sample. The Λ-coalescent has been developed as an alternative to Kingman’s coalescent to allow for multiple lineages to coalesce simultaneously [19]. This could prove useful for densely sampled infectious disease outbreaks in which multiple mergers are more likely to be observed, though remains to be generalised to arbitrary offspring distributions.

## Acknowledgements

We thank Erik Volz, Nicholas Grassly and Patrick Hoscheit for useful discussions. We acknowledge funding from the MRC Centre for Outbreak Analysis and Modelling at Imperial College, and CF thanks the Li Ka Shing Foundation for funding at Oxford University. LML was supported by a studentship funded by the Medical Research Council [grant number G01360].

